## Supplementary Figures for "Synaptic and intrinsic membrane defects disrupt early neural network dynamics in Down syndrome"

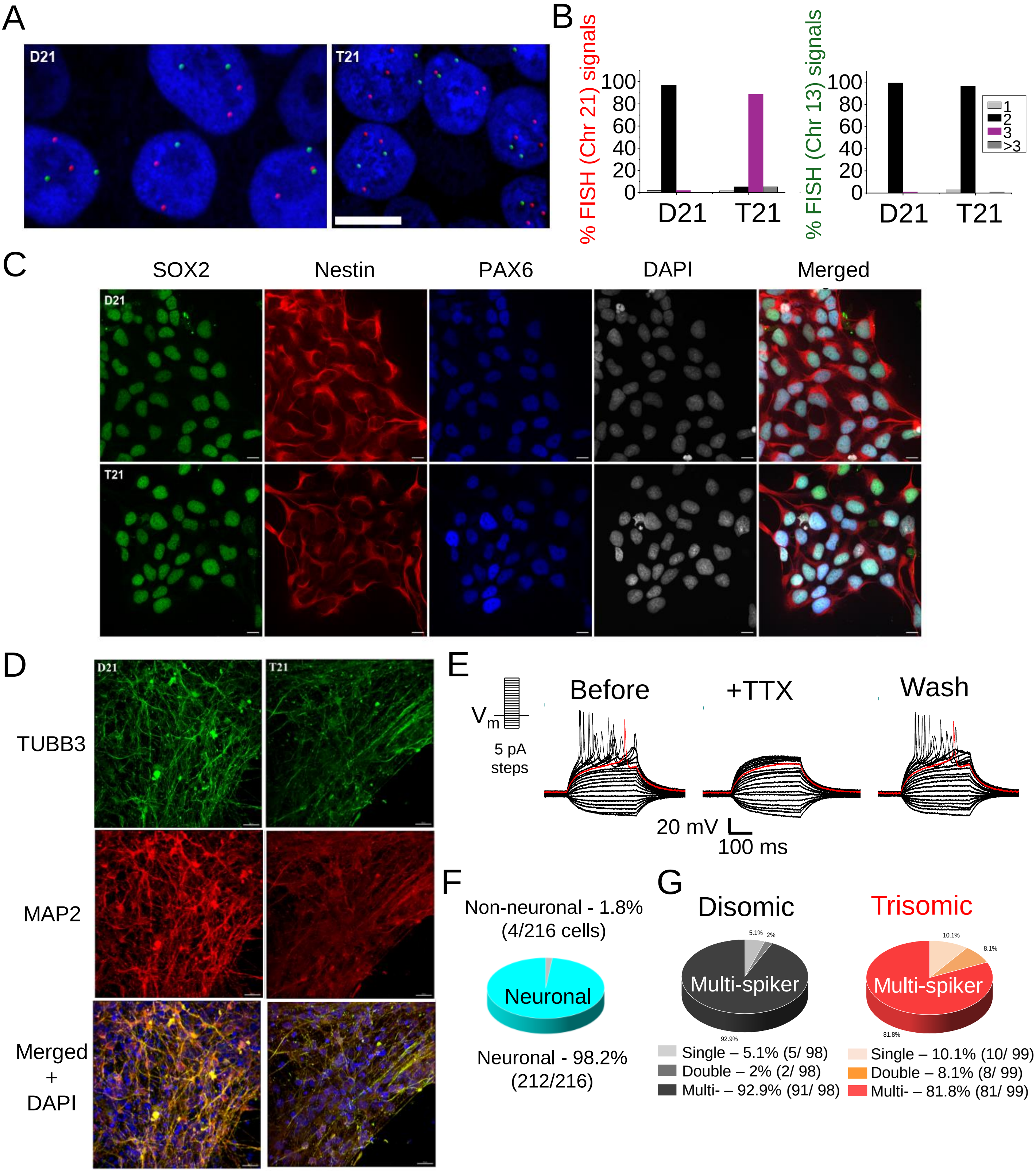

A
D21
T21
B
% FISH (Chr 21) signals
100
80
60
40
20
0
D21
T21
% FISH (Chr 13) signals
1
2
3
>3
C
SOX2
Nestin
PAX6
DAPI
Merged
D
TUBB3
MAP2
Merged + DAPI
E
Before
+TTX
Wash
Vm
5 pA steps
20 mV
100 ms
F
Non-neuronal - 1.8% (4/216 cells)
Neuronal
Neuronal - 98.2% (212/216)
G
Disomic
Trisomic
Multi-spiker
Single – 5.1% (5/ 98)
Double – 2% (2/ 98)
Multi- – 92.9% (91/ 98)
Single – 10.1% (10/ 99)
Double – 8.1% (8/ 99)
Multi- – 81.8% (81/ 99)

**Figure S1. – Validation of cell lines and neural differentiation**

*A,* Confocal images of disomic (D21) and trisomic (T21) neural stem cell nuclei (DAPI; blue) stained for chromosome 21 (red) using florescence in-situ hybridization. Chromosome 13 (green) staining has been used as a control. Scale bars - 10 µm. *B,* Quantification for chromosome 21 (left) and 13 (right) numbers in D21 and T21 cells showing presence of three copies of chromosome 21 in T21 cells. *C,* Representative images of isogenic disomic (top row) and trisomic (bottom row) cells stained with neural stem cell specific markers SOX2 (green), Nestin (red), PAX6 (blue) and counterstained with DAPI (gray). Scale bars - 10 µm. *D,* Representative images of mature isogenic disomic (D21, left column) and trisomic (T21, right column) neurons stained with pan-neuronal markers TUBB3 (green) MAP2 (red) and counterstained with DAPI (blue). Scale bars - 20 µm. *E,* Action potentials elicited by injection of constant current steps in human Down syndrome neurons. The current at which the neuron fires the first action potential (rheobase) has been depicted in red. Application of tetrodotoxin (TTX; 0.5 µM) abolishes action potentials which can be recovered upon wash off. *F,* Percentage of cells that fire stereotypical action potentials in whole cell current clamp recordings. 114 disomic and 102 trisomic cells from seven batches were recorded in current clamp and overall >98% of cells characterized using single cell electrophysiology were neurons based on action potential firing ability. Note that single cell electrophysiology relies on selecting cells using transmitted light microscopy factoring in morphology, quality of membrane and cellular health. *G,* Proportion of disomic and trisomic neurons that fired a maximum of one spike, two spikes or more than two spikes (multi-spiker). After establishing current clamp and recording basal spiking, a step current injection protocol was initiated allowing for saturation of spike input-output curve enabling the determination of maximum spiking properties.

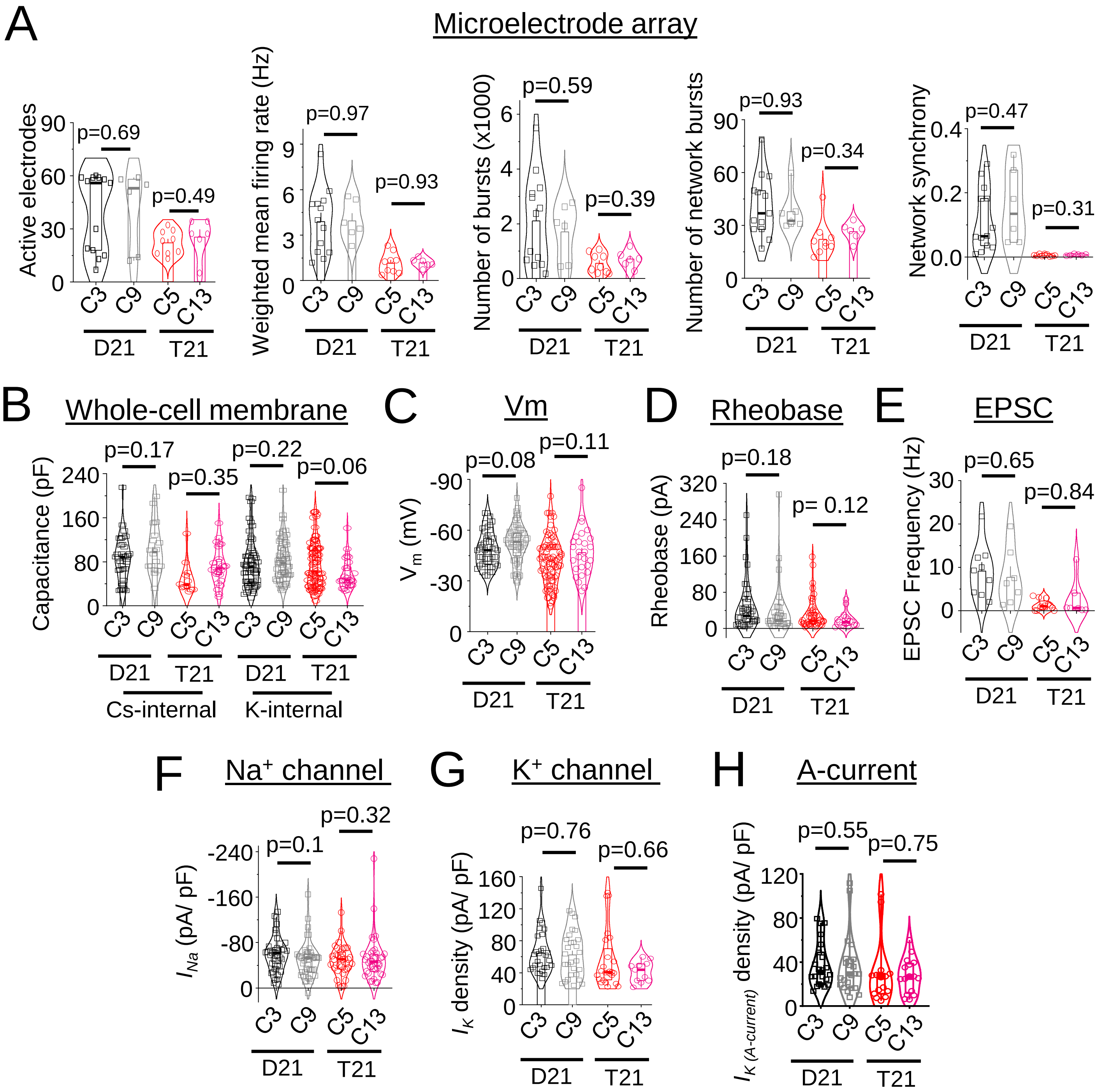


**Figure S2. Absence of overt inter-clonal functional variations within disomic (C3/ C9) and trisomic (C5/ C13) iPSC lines**

***A***, Number of active electrodes, weighted mean firing rate, number of electrode bursts, number of network bursts and synchrony index of C3 and C9 disomic lines along with C5 and C13 trisomic lines in microelectrode array recordings of weeks 9-10 iPSC-derived neurons. ***B***, Whole cell membrane capacitance of disomic and trisomic clones recorded using a Cs- or K-based internal solution. ***C-H***, Resting membrane potential ($V_m$) *(C)*, rheobase for spike firing *(D)*, frequency of excitatory postsynaptic currents (EPSCs) in a zero $Mg^{2+}$ bath solution *(E)*, $Na^+$ channel current density using a depolarizing single step protocol *(F)*, steady-state $K^+$ channel current density at a 90 mV step of I-V curve *(G)*, and A-type $K^+$ channel current density *(H)* of C3 and C9 disomic lines along with C5 and C13 trisomic lines. n = 6-13 wells in *(A)*, n = 7-87 cells in *(C-H)*; two tailed-unpaired t-test or Mann-Whitney test between disomic or trisomic clones.

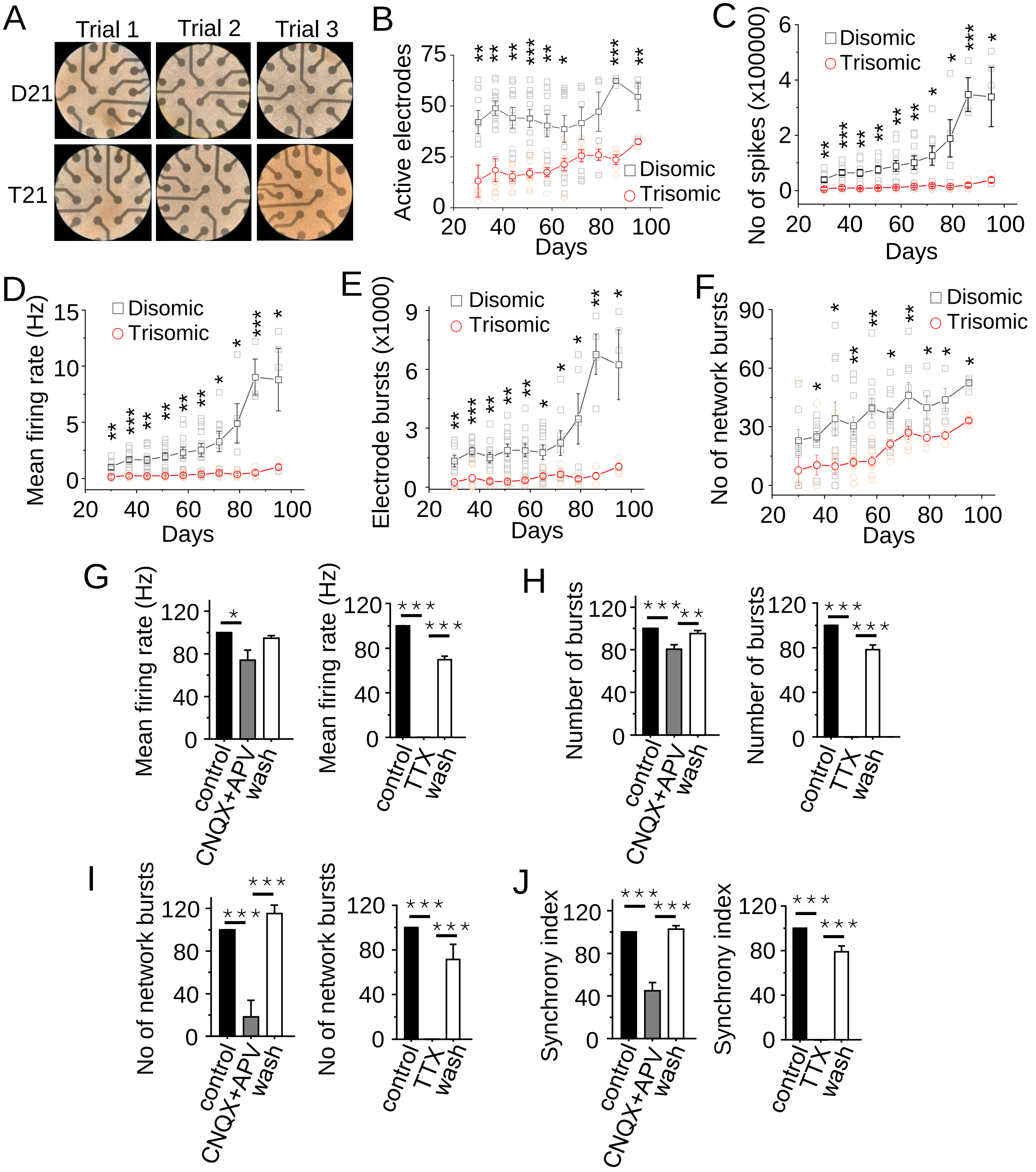
A
Trial 1
Trial 2
Trial 3
D21
T21
B
Active electrodes
0
25
50
75
20
40
60
80
100
Days
Disomic
Trisomic
C
No of spikes (x100000)
0
2
4
6
Days
Disomic
Trisomic
D
Mean firing rate (Hz)
0
5
10
15
Days
Disomic
Trisomic
E
Electrode bursts (x1000)
0
3
6
9
Days
Disomic
Trisomic
F
No of network bursts
0
30
60
90
Days
Disomic
Trisomic
G
Mean firing rate (Hz)
0
40
80
120
control
CNQX+APV
wash
Mean firing rate (Hz)
control
TTX
wash
H
Number of bursts
control
CNQX+APV
wash
Number of bursts
control
TTX
wash
I
No of network bursts
control
CNQX+APV
wash
No of network bursts
control
TTX
wash
J
Synchrony index
control
CNQX+APV
wash
Synchrony index
control
TTX
wash

**Figure S3. Neural network defects due to trisomy of chromosome 21**

*A,* Transmitted light images of microelectrode array recording electrodes with cells growing on top of them at 14 weeks from three different disomic (D21; top row) and trisomic (T21; bottom row) trial wells. Reduced activity of trisomic cells is not due to absence of cells in contact with recording electrodes. *B,* Number of active electrodes in 64-electrode wells during development. Trisomic cells consistently have fewer active electrodes suggesting reduced spiking activity in these networks. *C,* Number of total spikes recorded across development. The number of spikes rises rapidly as disomic cells mature but trisomic cells lag behind at all ages studied. *D,* Mean firing rate of disomic and trisomic cells across development and here too trisomic cells lag behind consistently. *E,* Number of electrode bursts across development showing deficits in bursting properties of trisomic neurons. *F,* Number of network bursts across development showing reduced network bursting of trisomic neurons. *G-J*, Normalised mean spike firing rate *(G),* number of electrode bursts *(H)*, number of network bursts *(I)* and synchrony index *(J)* of iPSC-derived disomic and trisomic neurons in presence of the AMPA receptor antagonist CNQX (10 µM) and the NMDA receptor antagonist APV (25 µM) or the $Na^+$ channel blocker tetrodotoxin (0.5 µM; TTX). Data has been normalised to basal recordings immediately prior to the application of the pharmacological agents. Note a silencing of spiking and bursting in TTX and reduction of single electrode and network bursts in CNQX and APV confirming that glutamatergic synaptic and intrinsic membrane properties determine spiking and network dynamics in these cortical networks. Results in *A-F* are from five different differentiations of two different lines and at least four wells per time point. In *G-J*, n = 6-8 wells. *P<0.05, **P<0.01. ***P<0.0001, Mann-Whitney test, two-tailed unpaired t-test between or one-way ANOVA.

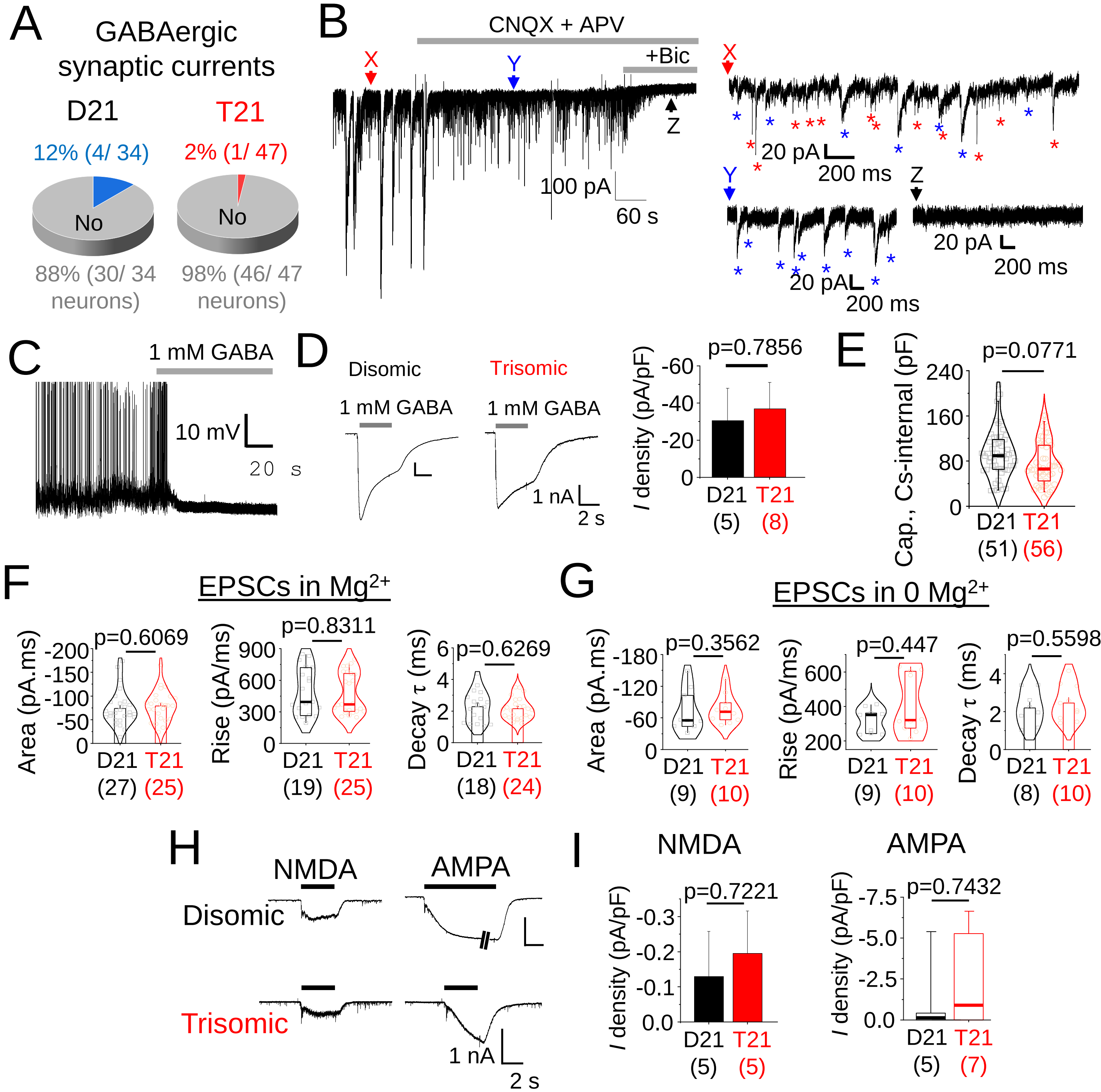

A
GABAergic synaptic currents
D21
T21
12% (4/ 34)
2% (1/ 47)
No
No
88% (30/ 34 neurons)
98% (46/ 47 neurons)
B
CNQX + APV
+Bic
X
Y
Z
100 pA
60 s
20 pA
200 ms
C
1 mM GABA
10 mV
20 s
D
Disomic
Trisomic
1 mM GABA
1 nA
2 s
p=0.7856
I density (pA/pF)
D21 (5)
T21 (8)
E
p=0.0771
Cap., Cs-internal (pF)
D21 (51)
T21 (56)
F
EPSCs in Mg2+
p=0.6069
Area (pA.ms)
D21 (27)
T21 (25)
p=0.8311
Rise (pA/ms)
D21 (19)
T21 (25)
p=0.6269
Decay τ (ms)
D21 (18)
T21 (24)
G
EPSCs in 0 Mg2+
p=0.3562
Area (pA.ms)
D21 (9)
T21 (10)
p=0.447
Rise (pA/ms)
D21 (9)
T21 (10)
p=0.5598
Decay τ (ms)
D21 (8)
T21 (10)
H
NMDA
AMPA
Disomic
Trisomic
1 nA
2 s
I
NMDA
AMPA
p=0.7221
I density (pA/pF)
D21 (5)
T21 (5)
p=0.7432
I density (pA/pF)
D21 (5)
T21 (7)

**Figure S4. GABAergic postsynaptic currents, EPSC kinetics and AMPA and NMDA whole cell currents**

*A,* Pie chart showing the fraction of cells that had GABAergic inhibitory postsynaptic currents (IPSCs) in cortical glutamatergic iPSC neurons. A limited number of cells (12% disomic (D21) and 2% trisomic (T21)) cells from only one differentiation out of eight received GABAergic inputs. The results from this differentiation have not been incorporated in the analysis. *B,* An example trace showing GABAergic postsynaptic currents of an excluded disomic cell. When present, GABAergic synaptic currents were identified by their slower kinetics (blue asterisk) compared to excitatory postsynaptic currents (red asterisk) in addition to application of the $GABA_A$ receptor antagonist bicuculline (25 µm; +Bic). Section X (red arrow) in the absence antagonists contains a mix of excitatory and inhibitory postsynaptic currents. In Y (blue arrow), application of AMPA and NMDA receptor antagonists CNQX and APV abolishes excitatory events leaving GABAergic events. In Z (black arrow), bicuculline abolished inhibitory events confirming their GABAergic identity. *C,* Application of 1 mM GABA blocks spontaneous action potentials in an iPSC-derived disomic neuron showing the presence of functional $GABA_A$ receptor responses. This suggests that the absence of inhibitory postsynaptic currents in these cells are not due to the absence of $GABA_A$ receptors. *D,* Representative 1 mM GABA-activated currents and current densities of disomic and trisomic cells. *E,* Whole cell capacitance (Cap.), calculated from the area under a -10 mV step capacity discharge curve, shows a trend towards reduced sizes (but not statistically significant) of trisomic (T21) cells compared to their disomic (D21) counterparts. *F,* Unchanged charge transfer, rate of rise and decay times of excitatory postsynaptic currents (EPSCs) recorded in $Mg^{2+}$ containing saline solution. *G,* Similarly, EPSC kinetics of charge transfer, rate of rise and decay times are also unchanged in saline devoid of $Mg^{2+}$. *H,* NMDA- and AMPA-activated currents of disomic (D21) and trisomic cells (T21). NMDA (50 µM) was applied with the co-agonist glycine (10 µM) and AMPA (10 µM) was co-applied with cyclothiazide (50 µM) to prevent desensitization. Cells were held at a potential of -70 mV. *I,* Bar charts and box plots showing unchanged NMDA and AMPA current densities in disomic and trisomic iPSC-derived cortical glutamatergic neurons. n numbers shown in brackets under genotypes. n = 5-56 cells; two-tailed unpaired t-test and Mann-Whitney test

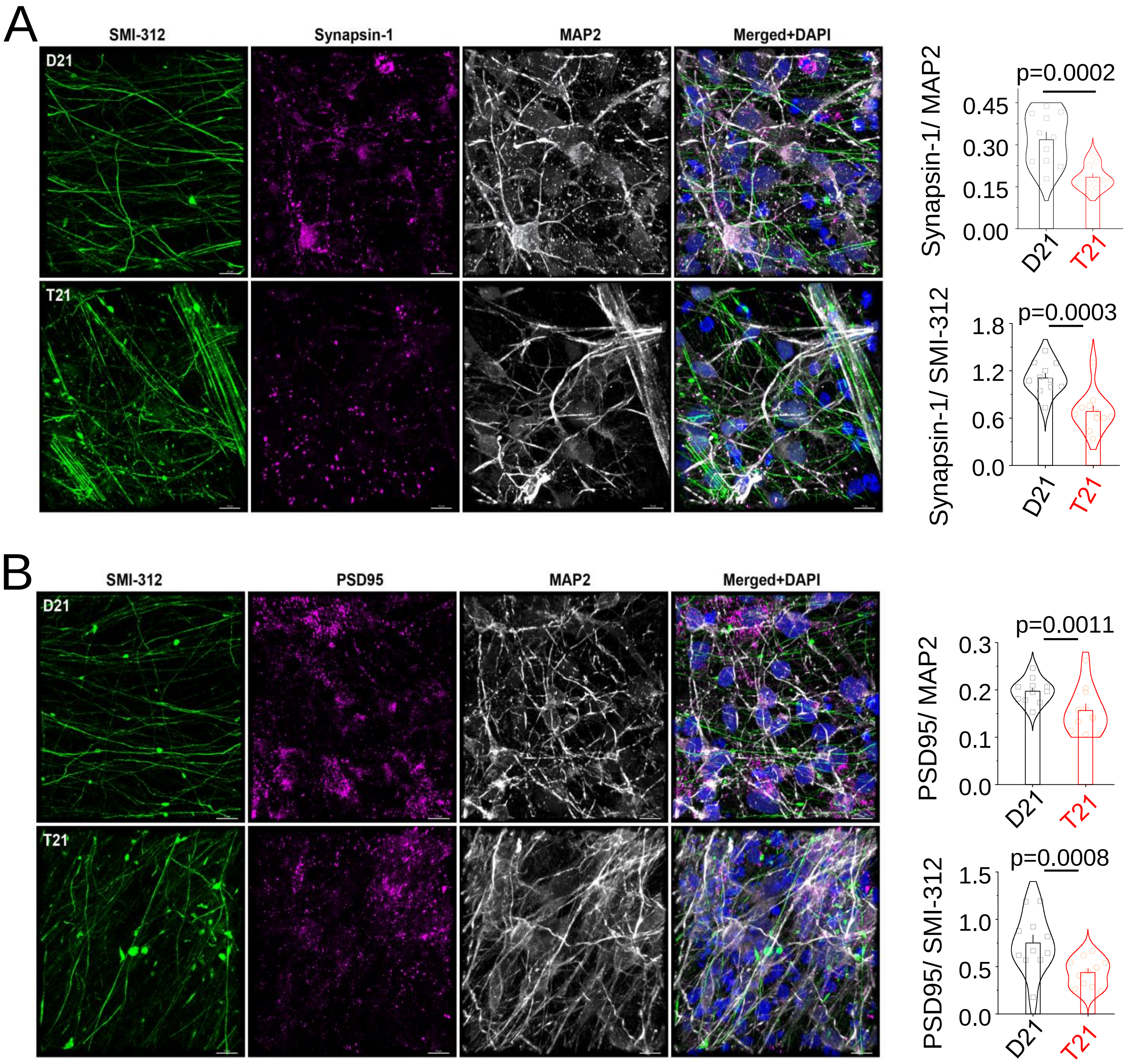


**Figure S5. Reduction of number of synapses in trisomy 21 neurons.**

*A, Left, r*epresentative images of mature isogenic neurons at sixty days *in vitro* stained with axon-specific marker (SMI-312; green), pre-synaptic marker synapsin-1 (magenta), dendrite-specific marker MAP2 (grey) and counterstained with DAPI (blue). *Right,* analysis showed significantly higher Synapsin-1 expression in both disomic (D21) dendrites and axons compared to trisomic (T21) cells. *B, Left,* Representative images of mature isogenic neurons at sixty days *in vitro* stained with axon-specific marker (SMI-312; green), post-synaptic marker PSD95 (magenta), dendrite-specific marker MAP2 (grey) and counterstained with DAPI (blue). *Right,* analysis showed significantly higher PSD95 expression in both D21 dendrites and axons compared to T21. Scale bars - 10 µm. Graphs represent means ± SEM. n = 10-11 3D image stacks. two-tailed unpaired t-test.

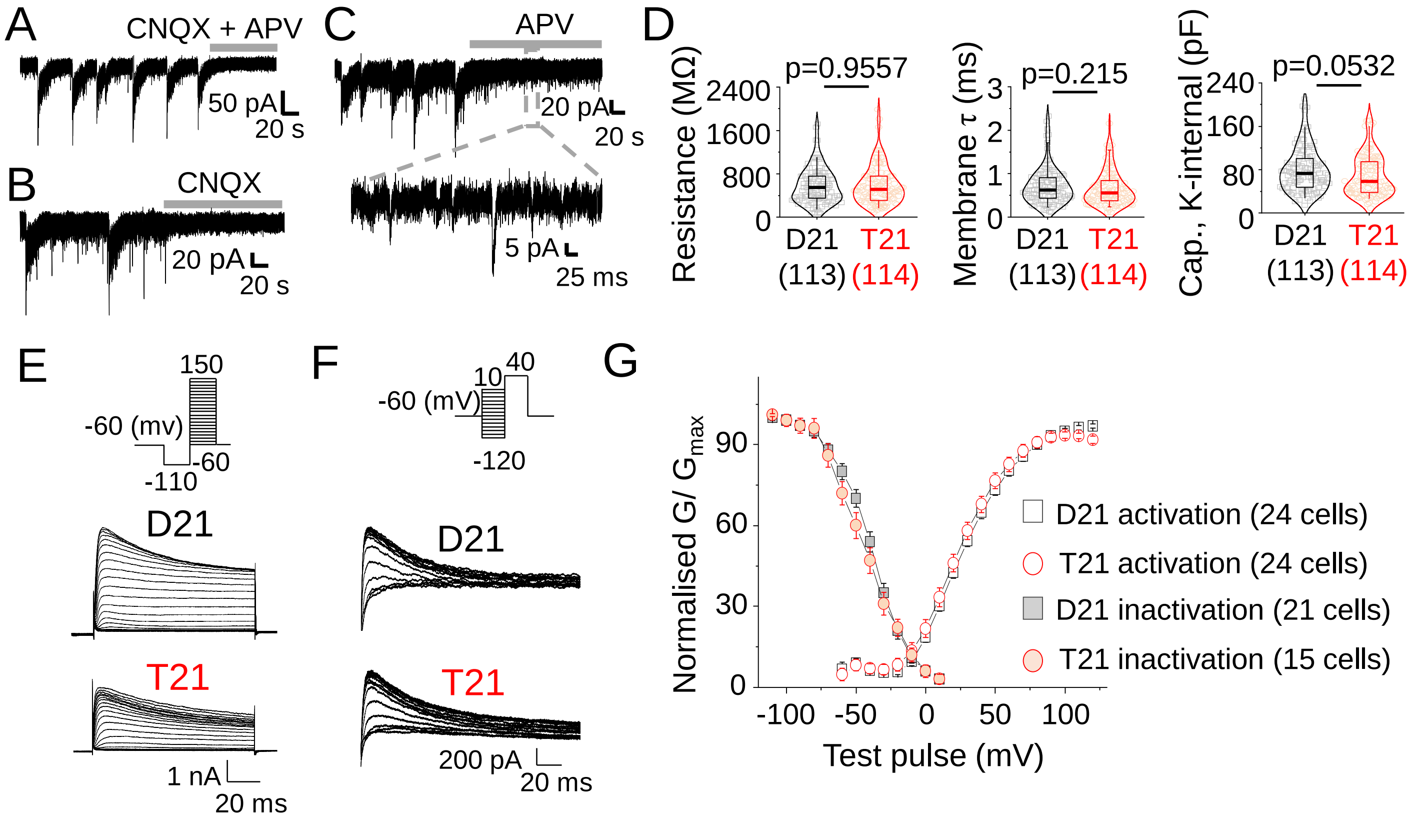


**Figure S6. Pharmacological properties of 0 $Mg^{2+}$ bursts, passive membrane properties and A-current activation and inactivation**

*A,* In 0 $Mg^{2+}$ saline, blocking AMPA and NMDA receptors with CNQX and APV abolishes bursts and excitatory postsynaptic currents (EPSCs) in our iPSC-derived cortical glutamatergic neurons. The example shown here is that of a disomic cell. Trisomic cells show similar block of bursting by CNQX and APV. *B,* Blocking AMPA receptors alone with CNQX also abolishes bursts and EPSCs suggesting the importance of AMPA receptor activity and EPSCs for bursting. *C,* Blocking NMDA receptors alone using APV can also abolish bursts but does not block EPSCs (inset) confirming NMDA receptor dependence of bursting. *D,* Input resistance and membrane time constant measured during current clamp recordings do not change between disomic (D21) and trisomic (T21) cells but the capacitance measured from the area under the curve using the same internal solution in response to a -10 mV hyperpolarising step trends towards smaller sized trisomic cells. *E,* Representative A-type $K^+$ channel currents recorded from disomic (D21) and trisomic (T21) neurons along with the protocol for activating these currents from a hyperpolarised potential (-120 mV) to more depolarized voltages. *F,* Representative A-type $K^+$ channel inactivation currents along with the step protocol. *G,* Normalized voltage-dependence of activation and inactivation profiles of A-type $K^+$ channel conductance does not change between disomic and trisomic neurons. N numbers of cells in brackets; Mann-Whitney test.

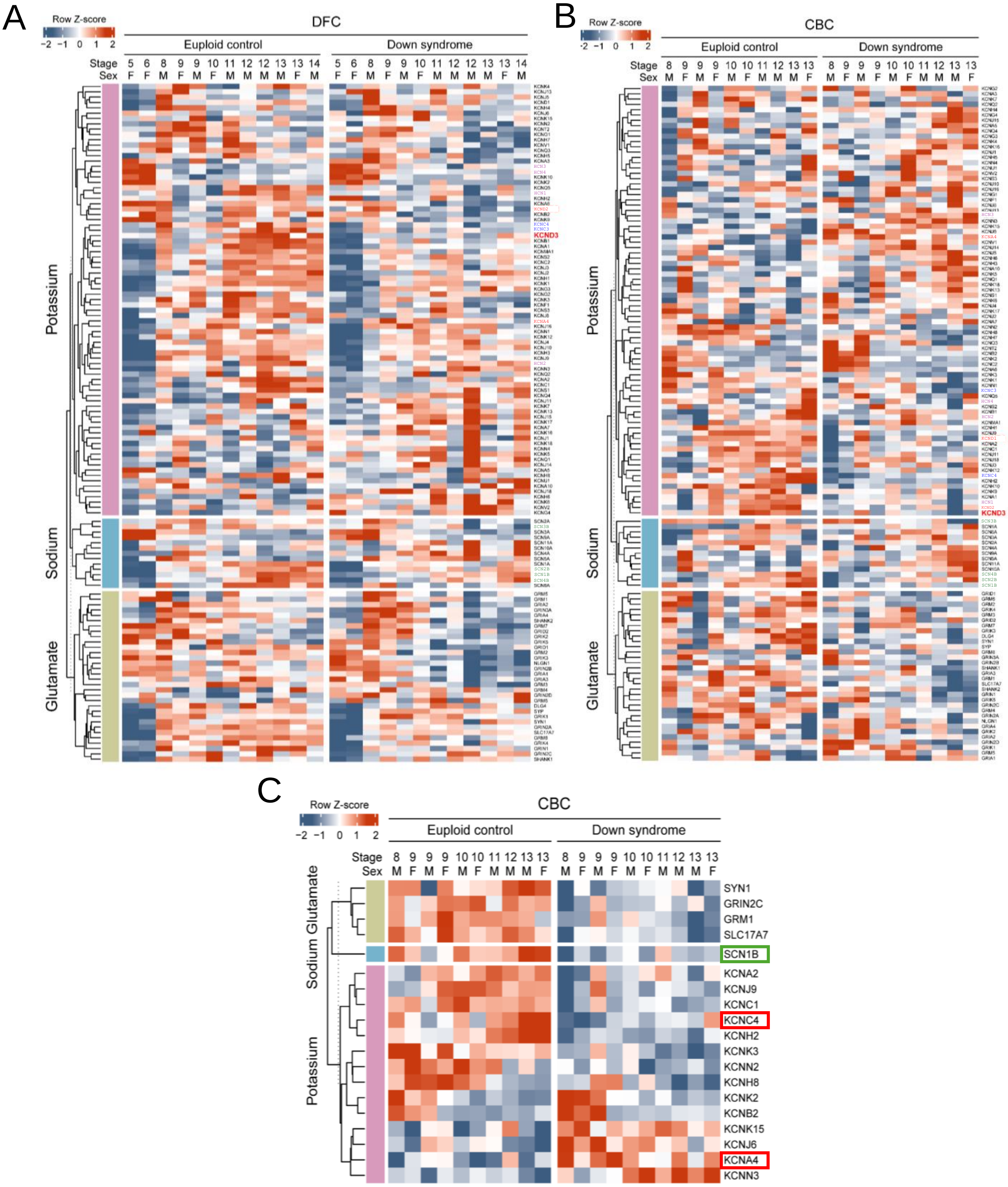
A
Row Z-score
-2 -1 0 1 2
DFC
Euploid control
Down syndrome
Stage
Sex
5 6 8 9 9 10 11 12 12 13 13 14
F F M F M F M M M M F M
5 6 8 9 9 10 11 12 12 13 13 14
F F M F M F M M M M F M
Potassium
Sodium
Glutamate
KCND3
B
Row Z-score
-2 -1 0 1 2
CBC
Euploid control
Down syndrome
Stage
Sex
8 9 9 9 10 10 11 12 13 13
M F M F M F M M M F
8 9 9 9 10 10 11 12 13 13
M F M F M F M M M F
Potassium
Sodium
Glutamate
KCND3
C
Row Z-score
-2 -1 0 1 2
CBC
Euploid control
Down syndrome
Stage
Sex
8 9 9 9 10 10 11 12 13 13
M F M F M F M M M F
8 9 9 9 10 10 11 12 13 13
M F M F M F M M M F
Glutamate
Sodium
Potassium
SYN1
GRIN2C
GRM1
SLC17A7
SCN1B
KCNA2
KCNJ9
KCNC1
KCNC4
KCNH2
KCNK3
KCNN2
KCNH8
KCNK2
KCNB2
KCNK15
KCNJ6
KCNA4
KCNN3

**Figure S7. Expression of $K^+$, $Na^+$ and HCN channels and glutamatergic synapse genes in the dorsolateral prefrontal cortex and cerebellar cortex at various stages of life from human brains**

*A-B,* Heatmaps showing expression of potassium channel-related and hyperpolarization-activated cyclic nucleotide-gated channel-related (purple), sodium channel-related (blue) and glutamate-related (green) genes in the dorsolateral prefrontal cortex (DFC) (A) or cerebellar cortex (CBC) (B) of euploid controls and Down syndrome brains. Note that HCN channels have been included in the list of $K^+$ channels. TEA-insensitive A-type $K^+$ channels have been highlighted with red text, whereas TEA-sensitive A-type $K^+$ channel genes have been depicted in blue text, HCN channel genes are in purple text and $Na^+$ channel β-subunit genes that affect channel kinetics have been depicted with green text. The z score indicates intensity of expression (red – high, blue – low). *C,* Heatmap showing changes of expression levels of potassium channel-related (purple), sodium channel-related (blue) and glutamate-related (green) genes in CBC in euploid controls and Down syndrome brains after birth. Changes to TEA-sensitive A-type $K^+$ channels have been highlighted with red boxes (*KCNC4* has mixed sensitivity depending on oligomerization states) and a $Na^+$ channel β-subunit gene that affects channel kinetics has been depicted with a green box. Note that *KCND3* is not a differentially expressed gene in this region. Table S1 contains a description of the stages of life.

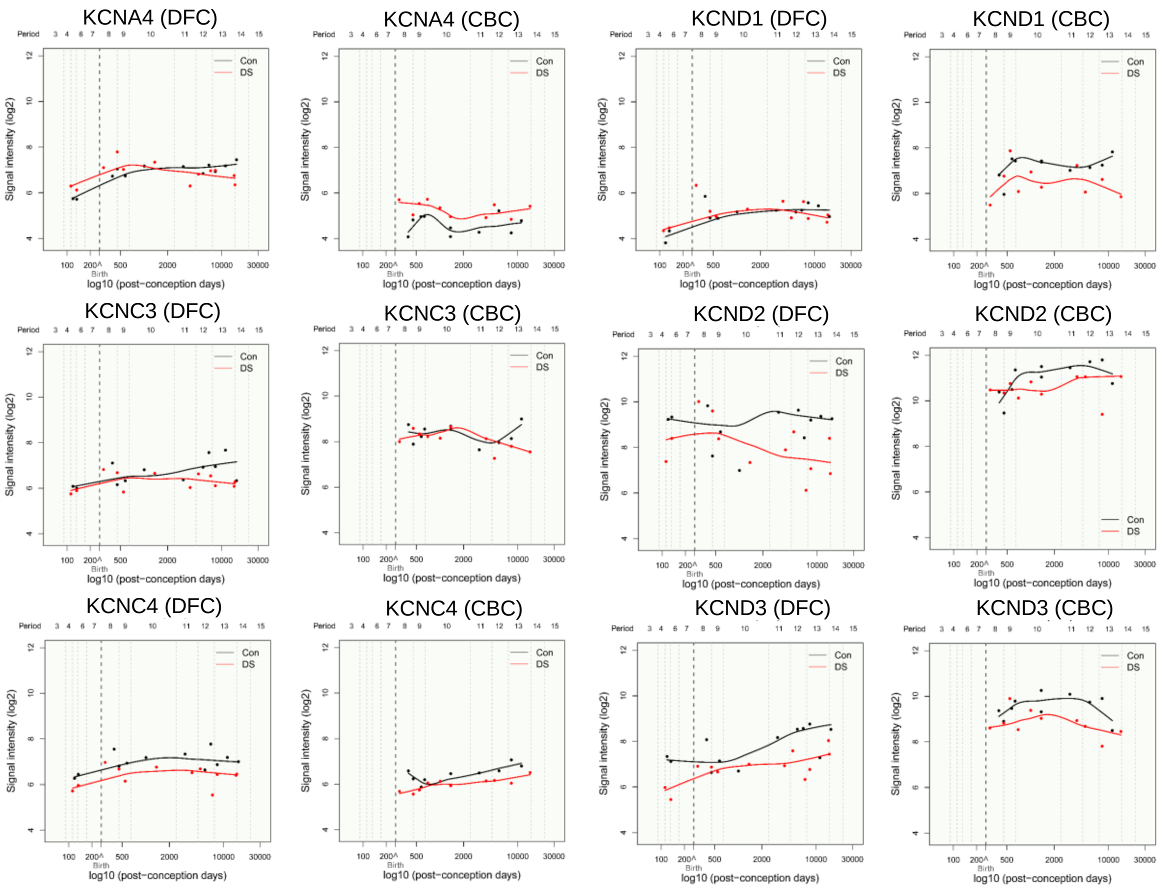


### Figure S8. A-current gene expression in dorsolateral prefrontal cortex and cerebellar cortex in Down syndrome

Gene expression patterns of *KCNA4*, *KCNC3*, *KCNC4*, *KCND1*, *KCND2* and *KCND3* are visualized using line graphs in the dorsolateral prefrontal cortex (DFC) and cerebellar cortex (CBC). Note that in the DFC, at early stages of development only expression of *KCND3* and *KCNC4* are significant. Table S1 contains a description of these stages. Stages 5 to 8 correspond to periods from mid-fetal development (16 weeks post-conception) to infancy (up to 6 months after birth).

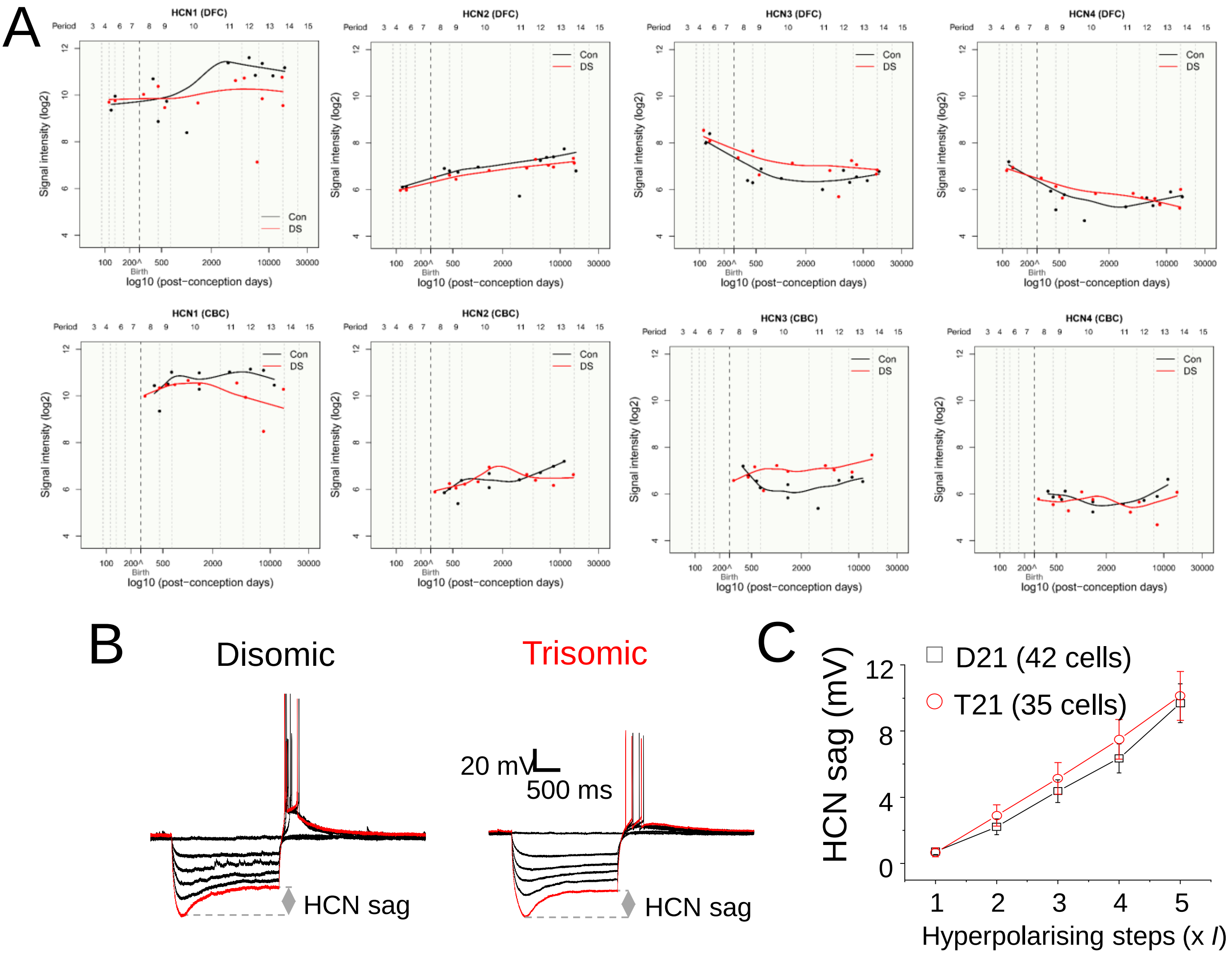


**Figure S9. Unchanged HCN gene expression and currents during early development in Down syndrome**

***A,*** Gene expression patterns of *HCN1*, *HCN2*, *HCN3* and *HCN4* visualized using line graphs are unchanged before birth in dorsolateral prefrontal cortex (DFC) and cerebellar cortex (CBC). ***B,*** Example HCN sags elicited upon hyperpolarisation of membrane voltage. Note rebound spikes upon release from hyperpolarization consistent with HCN activity. Hyperpolarising steps of currents were carefully injected in suitable equal increments such that the maximum hyperpolarising step yielded membrane potentials close to -150 mV or the cell underwent di-electric breakdown in which case the membrane voltage measures in the sweeps immediately prior to di-electric breakdown were analyzed. ***C,*** HCN sags do not change in trisomic (T21) cells compared to their disomic (D21) counterparts across a range of hyperpolarisations. n = 35-42 cells for HCN analysis, two-tailed unpaired t-test or Mann-Whitney test. Table S1 contains a description of the stages of life.
